# Oculomotor freezing indicates conscious detection free of decision bias

**DOI:** 10.1101/2021.08.19.456961

**Authors:** Alex L. White, James C. Moreland, Martin Rolfs

## Abstract

The appearance of a salient stimulus rapidly inhibits saccadic eye movements. Curiously, this “oculomotor freezing” reflex is triggered only by stimuli that the observer reports seeing. It remains unknown, however, if oculomotor freezing is linked to the observer’s *sensory experience*, or their *decision* that a stimulus was present. To dissociate between these possibilities, we manipulated decision criterion via monetary payoffs and stimulus probability in a detection task. These manipulations greatly shifted observers’ decision criteria but did not affect the degree to which microsaccades were inhibited by stimulus presence. Moreover, the link between oculomotor freezing and explicit reports of stimulus presence was stronger when the criterion was conservative rather than liberal. We conclude that the sensory threshold for oculomotor freezing is independent of decision bias. Provided that conscious experience is also unaffected by such bias, oculomotor freezing is an implicit indicator of sensory awareness.

**New & Noteworthy:** Sometimes a visual stimulus reaches awareness, and sometimes it does not. To understand why, we need objective, bias-free measures of awareness. We discovered that a reflexive freezing of small eye movements indicates when an observer detects a stimulus. Furthermore, when we biased observers’ decisions to report seeing the stimulus, the oculomotor reflex was unaltered. This suggests that the threshold for conscious perception is independent of the decision criterion and is revealed by oculomotor freezing.

## Introduction

You can often gain insight into another person’s mind by observing how they move their eyes and what they choose to look at. But even when they attempt to keep their gaze still, tiny involuntary eye movements reveal aspects of their mental state. Interspersed among slower types of fixational eye movements, involuntary *microsaccades* rapidly shift the gaze direction by small amounts (Rolfs 2009; Rucci and Poletti 2015). Microsaccades are in many ways similar to large saccadic eye movements (Hafed 2011; Otero-Millan et al. 2013; Rolfs et al. 2008), and their frequency and timing are affected by other cognitive and motor processes. For instance, microsaccade rates decrease in anticipation of sensory events(Abeles et al. 2020; Amit et al. 2019; Badde et al. 2020; Denison et al. 2019) and prior to voluntary eye and hand movements (Betta and Turatto 2006; Rolfs et al. 2006).

A particularly striking oculomotor phenomenon is oculomotor freezing (White and Rolfs 2016): saccadic eye movements are momentarily and automatically inhibited by the appearance of new stimuli(Engbert and Kliegl 2003; Hafed and Ignashchenkova 2013; Reingold and Stampe 2002; Rolfs et al. 2008). Specifically, the onset of a stimulus — be it auditory, tactile, or visual — causes a transient decrease in the spontaneous microsaccade rate that lasts from roughly 100 to 400 ms, which is followed by a brief rebound above baseline (Badde et al. 2020; Bonneh et al. 2015; Engbert and Kliegl 2003; Hafed and Ignashchenkova 2013; Rolfs et al. 2008; Scholes et al. 2015).

We recently found that oculomotor freezing is triggered only by stimuli that the observer detects (as measured by explicit report), revealing a possible link to visual awareness (White and Rolfs 2016). In those experiments, we presented brief grating stimuli (Gabor patches) on half the trials and asked the observers to report stimulus presence or absence. We developed an algorithm to convert microsaccade rates into a measure of oculomotor sensitivity (*o’*) that can be compared to perceptual sensitivity (*d’*). Contrast thresholds for the two sensitivity measures were indistinguishable (consistent with contemporaneous work by others (Bonneh et al. 2015; Scholes et al. 2015)). Crucially, the same physical stimulus gave rise to full-fledged oculomotor freezing when it was detected but caused no change in microsaccade rates when it was missed. Moreover, microsaccades were inhibited if observers reported having seen a stimulus even if none had appeared. Because of this correlation, a Bayesian algorithm could decode from observers’ eye movement patterns whether they had detected a stimulus or not. This oculomotor link to perception may provide a new tool for studies of perception in incommunicative patients, children, or non-human animals, and for “no-report” studies of consciousness (Tsuchiya et al. 2015).

The present study answers an important question left open by all previous studies: is oculomotor freezing triggered by observers’ *sensory experience*, or by their *decision* that a stimulus was present? Those two phenomena can be dissociated, and understanding which one lies at the origin of oculomotor freezing is vital to its interpretation and application. We consider two hypotheses to explain the established covariation between oculomotor responses and explicit perceptual detection (Bonneh et al. 2015; Denniss et al. 2018; Scholes et al. 2015; White and Rolfs 2016). Both assume a classical signal detection model: on each trial, the stimulus evokes an internal response that is compared against a criterion to decide whether to produce a response or not. Even when the physical stimulus and task demands are constant, the sensory response varies across trials, but the criterion is relatively stable. The two hypotheses concern whether the criterion for oculomotor freezing is the same as the criterion for explicit perceptual decisions.

1. *Shared criterion:* There is a single decision criterion that determines both explicit perceptual reports and oculomotor freezing. When the sensory response exceeds the criterion, it triggers both a “yes” decision and oculomotor freezing. The shared criterion can be strategically modified, to maximize expected rewards. In support of this possibility, manipulations of stimulus probability that affect decision bias affect activity in the superior colliculus (Crapse et al. 2018), which is also causally involved in controlling microsaccades (Hafed et al. 2009).
2. *Distinct criteria:* There are distinct criteria for triggering oculomotor freezing and for deciding that a stimulus was present. However, while the oculomotor criterion is inflexible, the observer can strategically change their perceptual decision criterion to maximize expected rewards as conditions change. Thus, the two criteria can diverge, breaking the link between explicit reports and oculomotor freezing. To explain our prior results(White and Rolfs 2016), this hypothesis assumes that the participants reported exactly what they perceived and set their decision criterion very near the criterion for oculomotor freezing.

We designed two experiments to discriminate between these hypotheses by manipulating observers’ decision criterion in a detection task. The first experiment used weighted payoffs (real money won or lost on each trial), and the second varied the expected probability that a stimulus would appear on each trial. Such manipulations shift the theoretically optimal criterion to a point that corresponds to a particular likelihood ratio *β_opt_* of target presence to absence, and have been shown to work empiricalally (Macmillan and Creelman 2005; Mulder et al. 2012; Swets et al. 1961). Our question here is whether and how these bias manipulations affect the prevalence of oculomotor freezing. To answer it, we conduct two main analyses of microsaccade rates: the first separates trials according to the physical stimulus presence, and the second additionally separates trials according to the participants’ reports of stimulus presence or absence. The shared-criteria hypotheses predict an effect of bias condition in the first analysis but not the second, the distinct-criteria hypothesis predicts the opposite.

## METHODS

Both experiments were pre-registered (https://osf.io/ycjgr; https://osf.io/s9myc/). The Ethics Committee of the German Society for Psychology (DGPs) approved the study.

### Experiment 1

#### Participants

We recruited a total of 16 observers from the Humboldt-Universität zu Berlin community, with normal or corrected-to-normal vision. They participated in exchange for a payment that depended on performance (details below). Of the 14 observers who completed the study (see below), 6 were male, 8 were female, and their ages ranged from 19 to 34 years (mean 26.3). All were naive as to the research aims, and gave informed consent.

The sample size was chosen on the basis of a power analysis based on the data from White & Rolfs (2016). In Experiment 3 of that study, we found an effect of orientation adaptation on microsaccade rates. That effect size was modest: the maximal difference at 350 ms post-stimulus was 0.2 Hz. Averaging over the time window when the overall inhibitory effect of stimulus presence was significant, the mean effect was 0.13 Hz.

We made the conservative assumption that if there is an effect of payoff condition, it is 75% as large as the effect of orientation adaptation, at each individual timepoint. We conducted a power analysis to determine how many participants would be necessary to find such an effect with a power of 0.8. For each possible sample size (N) between 10 and 20, we simulated 100 experiments. For each experiment, we conducted a bootstrapping analysis: in each of 1000 repetition, we drew N observers with replacement from the original data set in White & Rolfs (2016). For each observer, we computed the difference in microsaccade rate between the unadapted and adapted condition, at each time point post-stimulus, multiplied by 0.75. We then averaged those differences across the resampled participants. Over 1000 repetitions we built up a distribution of differences at each time-point, from which we could extract a p-value. We applied the false discovery rate correction to determine at which time-points the difference was significant by applying. For each simulated experiment, we considered the overall effect to be significant if the difference was significant in at least 10 individual time-points. For each N, we defined power as the proportion of experiments with a significant effect. The minimal N to have a power over 0.8 was 14 (estimated power = 0.87).

Two participants began the study but did not finish it and were not included in the analyses. One was unable to finish all the sessions, and another discontinued after three sessions with d’ far above the acceptable range, due to threshold estimation failure. Thus, the final sample included 14 participants.

#### Apparatus and Stimuli

Observers sat in a darkened room with their head on a chin rest, 270 cm from a projection screen that displayed stimuli with a gamma-linearized ProPixx projector (VPixx Technologies; 120 Hz, 1920 x 1080 pixel resolution). We recorded the gaze position of both eyes at 500 Hz with a head-mounted Eyelink 2 system (SR Research, Ontario, Canada). Stimuli were controlled and data collected with the Psychophysics and Eyelink toolboxes (Brainard 1997; Cornelissen et al. 2002; Pelli 1997). The grayscale display (1920 x 1080 pixels, 120 Hz refresh rate) had 8 bits of resolution in luminance. The background luminance was set to 35% of its maximum (18.15 cd/m^2^).

The fixation mark was a 4 by 4-pixel black-and-white checkerboard pattern of width 0.09 degrees of visual angle (dva). In between trials, this mark was replaced by a circle (0.27 dva radius) of alternating black and white pixels. The target stimulus was a Gabor pattern: a 0.75 cycles/dva, vertically oriented sinusoidal grating windowed by a two-dimensional Gaussian (σ = 0.67 dva).

#### Procedure

Observers began each trial by fixating on the central mark. After 0.5–2.5 s, the target Gabor stimulus flashed for 8.3 ms. The target’s onset time had a roughly flat hazard rate: on each trial, the onset time was set to 0.5 s plus a value drawn from an exponential distribution (Mean = 0.65 s) clipped at 2 s. The target’s phase on each trial was randomly set to either 0° or 180°. On 50% of the trials, the target had non-zero contrast (target-present trials). On the remaining trials, its contrast was set to 0, causing no change on the screen (target-absent trials). The fixation mark remained visible at the center of the Gabor. 492 ms after target onset, a beep (400 Hz, 50 ms, delivered through headphones) indicated that the trial was over.

The observer’s task was to indicate whether the target was present or absent by pressing the up or down arrow, respectively, with the right hand. Response time was unlimited, but responses were not allowed before the beep. Tones delivered immediately after the response indicated whether the response was correct or incorrect, and how many points were won or lost (details in the next section). After an inter-trial interval (700 ms) containing only the circular fixation mark, the next trial began.

The first session began with practice and then two blocks of staircase trials to estimate the observer’s contrast threshold. During the staircase blocks, the contrast was adjusted after each trial according to the single-interval adjustment matrix (SIAM) staircase procedure (Kaernbach 1990). The contrast adjustment depended on the stimulus and response: after a hit, −0.3 log_10_ units; miss, +0.3 log_10_ units; false alarm, +0.6 log_10_ units; correct rejection, no adjustment. The magnitudes of these steps were halved after the 1st and 2nd staircase reversals. In each block, we interleaved two staircases, one starting at a relatively high and the other at a low level of contrast. The block ended when both staircases underwent 10 additional reversals. The mean contrast of all but the first 2 reversal points provided the threshold estimate. We defined the observer’s contrast threshold as the mean of 4 threshold estimates (2 from each of 2 blocks).

In the main experimental blocks (80 trials each), the target’s contrast was set to the observer’s estimated threshold. The mean stimulus contrast in included trials was 9% (ranging across individuals from 7% to 12%).

#### Payoff conditions

Our main manipulation is to the reward structure for correct and incorrect responses on target-present and target-absent trials. On each trial the observer won or “points”, which at the end of the experiment were converted to a monetary payment (1600 points = €1). By varying payoffs, we aimed to manipulate the observer’s *detection criterion:* that is, how much internal sensory evidence is required for the participant to report “target present” (Macmillan and Creelman 2005; Swets et al. 1961). In the main experimental blocks, there were two payoff conditions: conservative and liberal. Additionally, a neutral condition was used in the initial staircase blocks to estimate contrast threshold. Following classic signal detection theory, we assumed that on each trial the observer bases their decision on a single value *r*, which is the amount of sensory evidence in favor of target presence. The probability distribution of *r* on target-absent trials is *f_a_*(*r*), a Gaussian with *μ*=0 and *σ*=1. The probability distribution of *r* on target-present trials is *f_p_*(*r*), a Gaussian with *μ*=*d’* and *σ*=1. *d’* is the observer’s sensitivity to the target. The observer’s criterion can be expressed as *c*, the cutoff value of *r* needed to report presence. A related measure is the observer’s bias, the likelihood ratio *β:*

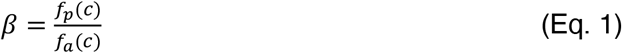

After substituting the full Gaussian formulas for *f_p_* and *f_a_*, we can reduce the equation to:

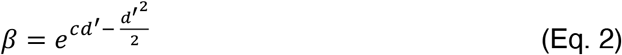

The payoffs in each condition were set to achieve a desired optimal criterion *β_opt_*: the value of *β* that maximizes the expected reward. The values of *β_opt_* were 3 for the conservative condition, 1 for the neutral condition, and 1/3 for the liberal condition. We set the payoffs such that the optimal observer, with a *d’* of 1.5, would earn an average of 6.4 points per trial. Over 1280 trials, that would yield a payment of €5.12 at our exchange rate of 1600 points/€. By setting the target luminance contrast to detection threshold, we aimed to keep each observer’s *d’* near 1.5. Given the average expected reward/trial (6.4 points) and the expected d’, we computed the payoff matrix that would lead an ideal observer to set their criterion to the desired *β_opt_*. Specifically, we computed the payoffs for target-present trials, *R_p_*, and for target-absent trials, *R_a_*. For each trial type *j* (*j*=*p* for target-present; *j*=*a* for target-absent), the reward for correct responses is *R_j_* points and the reward for errors is –*R_j_* points.

On any given trial, there were four possible outcomes: hits or misses if a target was present, or correct rejections or false alarms if there was no target. Given *d’* and *β*, we can compute the probabilities of each of those outcomes. Given *R_p_* and *R_a_*, we can then compute the expected reward *V* per trial:

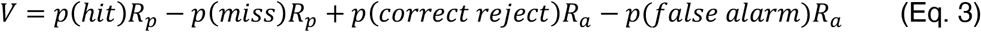

Given that the prior probabilities of target presence and absence were both equal to 0.5, the optimal likelihood ratio criterion is the ratio of payoffs:

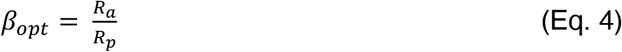

Therefore, greater payoffs on target-absent trials should induce a conservative (higher) criterion, whereas greater payoffs on target-present trials should induce a liberal (lower) criterion. In our conservative condition (*β_opt_* = 3), payoffs on target-absent trials should be three times payoffs on target-present trials. The inverse is true in the liberal condition. Working backwards from the equations above, and given our desired *d’* and expected reward per trial (*V*), we computed the payoff matrix shown in **Table 1**.

**Table 1:** Payoff matrix. For each condition, this table lists the number of points that can be won (positive values) or lost (negative values) for each type of response. The neutral condition was only used in the initial staircase blocks.

| Condition | Hit ($R_p$) | Miss ($-R_p$) | Correct reject ($R_a$) | False alarm ($-R_a$) |
| --- | --- | --- | --- | --- |
| Conservative | 4.9 | -4.9 | 14.8 | -14.8 |
| Liberal | 14.8 | -14.8 | 4.9 | -4.9 |
| Neutral | 11.7 | -11.7 | 11.7 | -11.7 |

The payoff on each trial was indicated by a feedback tone immediately after the response. These tones were composed of one, two, or three beeps, depending on the absolute value of the payoff (as shown in **Table 1**, there were three possible magnitudes). When there were multiple beeps, their pitches ascended in a major scale for correct responses or descended in a minor scale for incorrect responses. Each beep was separated by 20 ms of silence. In the liberal condition, for example, hits won 14.8 points and were followed by three ascending beeps, whereas false alarms cost 4.9 points and were followed by one low-pitched beep. The three beeps used for correct tones were: 75 ms of 440 Hz; 80 ms of 587 Hz; and 85 ms of 659 Hz. The three beeps used for incorrect feedback tones were: 75 ms of 196 Hz; 80 ms of 155 Hz; and 85 ms of 131 Hz.

The total number of points won were displayed at the end of each block. Prior to each block, instructions regarding the payoff structure were displayed on the screen. These instructions consisted of a 2×2 table showing the number of points that could be won or lost for reporting “Yes” or “No” depending on whether a target was present or absent. The values in this table were the same as in the corresponding condition’s row in **Table 1**, rounded to the nearest integer. A single sentence was written above the table: in the Conservative condition, “Rewards and penalties are greatest when the target is absent.”; in the Liberal condition, “Rewards and penalties are greatest when the target is present.”

Importantly, the words “liberal” or “conservative” were never said to the participants, nor did experimenters tell them what the optimal strategy was for any given condition. However, in the first training session, the participant read a longer document of instructions that said, “In the main part of the experiment, we will vary the number of points you can win or lose depending on presence of the target and the response you make. There are two types of blocks that differ in the relative rewards and penalties on trials when the target was really present or absent. To win the most money, you should adjust how sure you need to be to say ‘yes’ or ‘no’, depending on the points available for each type of response in the current block.” When introducing the conservative condition, the instructions said: “You will win <u>three times</u> as many points when the target is absent and you say no, than when a target is present and you say yes…and lose <u>three times</u> as many points when the target is absent and you say yes, than when a target is present and you say no.” Complementary instructions followed for the liberal condition. Observers were also instructed that they could win points and earn money during the staircase blocks as well as the main blocks.

In the first session, we informed observers that they would be paid a base hourly rate of €7/hour, plus a bonus equal to the total number of points they accumulated during the trials, divided by 1600. The maximum bonus they could earn in any given hour-long session was €4. The mean bonus paid for two main experimental sessions was €4.66 (range €4.13 to €5.30).

Each participant completed a total of 8 blocks of each condition (for a total of 640 trials/condition). The first session began with practice, the staircase to estimate threshold, and if time permitted, some main experimental blocks. In each subsequent session (about one hour each), the typical observer completed 8 blocks: the first four of one payoff condition, and the next four of the other condition. In each session, observers thus did an equal number of blocks of the two payoff conditions. The order of conditions alternated across sessions, and a random half of the observers started with the liberal condition.

Completing all 16 blocks required a total of three sessions for the typical participant (including the first staircase session). At the start of the 2^nd^ and 3^rd^ sessions, a practice block established whether the prior session’s contrast threshold was still appropriate; in some cases, it was necessary to re-evaluate the threshold and re-set the contrast level for that session to keep *d’* near 1.5. If the overall *d’* in a full session (~8 blocks) was above 2.0 or below 1.0, we excluded those blocks from analysis and re-ran them in an extra session. This occurred when our threshold estimate was significantly inaccurate. A total of three sessions from three participants were excluded and re-run in that fashion. The reason to exclude them is that our analyses of interest depend on the target stimulus being at threshold visibility. Importantly, we always excluded and re-ran the same number of blocks of each payoff condition.

#### Eye-tracking

At the start of each block, we performed a 9-point calibration within a central square region, 21 dva wide. Every 28 trials, we performed a standard drift correction by having the observer press a key while fixating a dot at the screen’s center. If either eye’s gaze position deviated more than 2 dva from the fixation mark between the start of a trial start and the beep, that trial was immediately terminated and repeated at the end of the block. We also detected fixation breaks offline by defining, for each trial, the fixation position as the median gaze coordinates during the first 100 ms of the trial, and fixation breaks as deviations >2 dva from that. Trials with offline-detected fixation breaks were excluded from the analysis, but that only excluded an average of 1 trial per participant (maximum 3).

### Experiment 2

#### Participants

We recruited a total of 20 observers from the Humboldt-Universität zu Berlin community. All had normal or corrected-to-normal vision, participated in exchange for payment, and gave informed consent. Of the 14 observers who completed the study and were included in the analysis (see below), 4 were male, 10 were female, and their ages ranged from 20 to 37 years (mean 25.4).

We used the same number of participants as in Experiment 1, but with twice as many trials per condition. The reason is that this experiment contained a condition in which the target was half as likely to appear (and we needed to separately analyze trials with and without targets). 6 participants were not included in the analysis because they discontinued participation before completing the study (in two cases because their d’ was out of range in one or more completed sessions and they declined to repeat them). Thus, the final sample included 14 observers.

#### Procedure

All stimuli and methods in Experiment 2 were the same as in Experiment 1, except as noted here. Observers began each trial by fixating on the central mark. Then a *probability cue* appeared for 1 s. The target probability cues were formed of 12 dots (each 0.2 dva in diameter) arranged in a ring around fixation (radius 3 dva). The dots on each trial were all of the same color, either cyan or magenta. For half the observers, a cyan cue indicated low target probability and magenta indicated high target probability. For the other half of observers, the colors were reversed. Then, after a variable delay of 0.5–2.5 s, the target Gabor stimulus flashed for 1 frame (8.3 ms), and the trial proceeded as in Experiment 1. The mean stimulus contrast in included trials was 6% (ranging across individuals from 5% to 9%).

#### Feedback and rewards

The feedback and reward structures were matched to the “neutral” condition in Experiment 1 (used in the staircase blocks). The participants won 11.7 points on correct trials (hits or correct rejections) and lost 11.7 points on incorrect trials (misses or false alarms). The feedback tones were two ascending beeps or two descending beeps.

#### Probability conditions

Our main manipulation was the probability of a target being present on each trial (*p_T_*). In “low-probability” trials, *p_T_* = 0.25, and on “high probability” trials, *p_T_* = 0.75. Those trials were randomly intermixed, because if they were in separate blocks, there could be hysteresis effects due to different amounts of stimulation in each block. The cyan or magenta pre-cue indicated the target probability condition at the start of each trial.

Given the average expected reward/trial (6.4 points) and the expected d’ (1.5), we computed the target probabilities that would lead an ideal observer to set their criterion to the desired *β_opt_*. Using the expected reward on each trial (Eq. 3), we can compute *β_opt_* from the ratio of payoffs, scaled by the ratio of the probability of no target and the probability of a target:

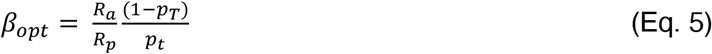

See Swets et al. (1961) for an equivalent derivation. In Experiment 2, *R_a_* = *R_p_* = 11.7 points. Therefore,

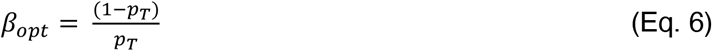

In the low-probability condition, *p_T_* = 0.25 and *β_opt_* = 3, the same as in the conservative payoff condition of Experiment 1. In the high-probability condition, *p_T_* = 0.75 and *β_opt_* = 1/3, the same as in the liberal payoff condition of Experiment 1. We therefore label the low-probability condition as the conservative condition, and the high-probability condition as the liberal condition.

At the start of the experiment, the observer was instructed to pay attention to the colored probability cues and was told their exact meaning. We did not tell the observers *how* to use the cues, but we did tell them, “If you pay attention to the colored dots and adjust your responses accordingly, you could gain roughly 20% more money than if you ignore them!”. Prior to each block, we displayed a reminder about what the probability cues mean.

Each participant completed a total of 32 blocks of the experiment (80 blocks per trial, for a total of 2560 trials, 1280 per condition). Completing all 32 blocks required a total of five or 6 sessions for the typical participant (including the first staircase session). The mean bonus paid for the main experimental sessions was €10.67 (range €7.61 to €13.64).

As in Experiment 1, we excluded and re-ran sessions with d’ above 2.0 or below 1.0. That occurred for a total of 5 sessions, one per each of 5 observers. On average, less than 0.1% of trials were excluded for offline fixation breaks (max 0.3%).

### Analyses

#### Perceptual data analysis

We excluded trials with reaction times >4 SDs above the observer’s median. Across participants, this criterion excluded an average of 1% of trials in Experiment 1 (maximum 1.6%), and an average of 0.7% in Experiment 2 (maximum 1.4%). We then computed perceptual sensitivity in each condition using the observer’s hit rate (HR, the proportion of ‘yes’ responses on target-present trials) and false alarm rate (FR, the proportion of ‘yes’ responses on target-absent trials):

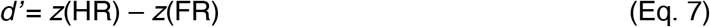

where *z* is the inverse of the normal cumulative distribution function. To avoid undefined *d’* values, HR and FR were not allowed to fall below 1/(2N) nor to exceed (1–1/(2N)), where N is the number of target-present or absent trials. For example, if the hit rate was 1, we assumed that, had we run twice as many trials, there would have been 1 miss. We also report the observer’s criterion

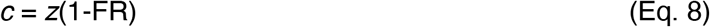

From that, we compute the bias *β*, the likelihood ratio, using Equation 2 defined above.

To evaluate the effect of payoff condition on these perceptual measures (*d’* and *β*), we used bootstrapping to estimate 95% confidence intervals between pairs of conditions. A difference is deemed significant if the 95% confidence interval excludes 0 (a two-tailed test).

#### Microsaccade detection

The trial exclusion criteria applied in the perceptual data analysis (see above) also applied to the eye movement analysis. Our analysis of eye movement traces followed the procedure reported in our previous paper (White and Rolfs 2016). We first transformed the raw gaze positions into velocities (dva/s) and smoothed them by averaging over neighboring pairs of two samples. Then, we identified microsaccadic events as shifts in gaze position with 2D velocities that exceed—for at least 3 samples—an ellipse with horizontal and vertical radii equal to five times the horizontal and vertical median-based standard deviations, respectively (Engbert and Mergenthaler 2006). However, for 6 observers in Experiment 1, and 3 in Experiment 2, the fixed threshold of 5 SDs yields very few microsaccades, so we lowered the threshold to 4.

Monocular microsaccadic events less than 10 ms apart were merged. We defined binocular microsaccades as those with at least 1 sample of overlap between the two eyes, and again, merged binocular microsaccades less than 10 ms apart. We defined microsaccade onset as the time the first of the two eye velocities exceeded the threshold, and offset as the timepoint just before the last eye’s velocity dropped below threshold. Other parameters (e.g., amplitude) were averaged over the two eyes. We included in the analysis only binocular microsaccades with durations ≥ 6 ms, amplitudes ≤ 1 dva, and peak velocities ≤ 250 dva/s.

#### Microsaccade rate analysis

We then determined the time-varying microsaccade rate for each experimental condition with a smoothing procedure. First, we counted the number of microsaccades detected at each millisecond t relative to target onset, across all trials in each condition. Then, for each time point t, we computed a weighted sum of microsaccades in the local interval, using a “causal” kernel:

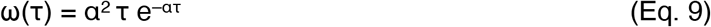

ω describes the weight given to microsaccades τ ms before time point t. We shifted the filter by 1/α ms to avoid a temporal bias and give the most weight to microsaccades at point t (Rolfs et al. 2008; Widmann et al. 2014). The parameter α was set to 1/25. The smoothed rate r(t) is the weighted sum of microsaccades divided by the total number of trials in the sample, and converted into Hz by multiplying by 1000. Microsaccade rates were computed from −350 to +500 ms relative to target onset.

To estimate the statistical significance of changes in microsaccade rates, we bootstrapped them by simulating 1000 repetitions of the experiment (Efron and Tibshirani 1993). On each repetition, we resampled with replacement from the set of observers then took the mean between conditions. That gave us distributions of differences at each time point. The two-tailed bootstrapped p-value is defined as twice the proportion of differences that fell below 0. When evaluating differences at many time points, we applied the false discovery rate correction (Benjamini and Hochberg 1995). Two conditions are deemed significantly different if the 95% confidence interval of differences does not include zero (corrected p<0.05). (Note: this bootstrapping procedure differs from what we pre-registered, in that it is simpler and focuses on variability across observers rather than variability across trials within each observer, thus being a nonparametric analogue of a t-test).

To directly compare changes in microsaccade rate to perceptual sensitivity, we computed an analogous estimate of oculomotor sensitivity (White and Rolfs 2016). At each millisecond, the lack of a microsaccade following a stimulus is a “hit”, and the lack of a microsaccade following no stimulus is a “false alarm”. From the resulting oculomotor hit rate (HR) and false alarm rates (FAR), we can compute oculomotor *d’_o_* at each time point t relative to stimulus onset (0<=t<=500):

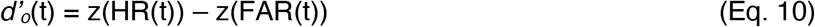

Like perceptual *d’*, this measure requires correction if HR or FAR reach extreme values. This can happen if no microsaccade were detected during a period around t as wide as the base B of the filter (~200 ms). Therefore, both rates will not be allowed to fall below 1/(2NB) nor to exceed (1–1/(2NB)), where N is the number of target-present or absent trials, respectively. That is, we assume that had we run twice as many trials, we would have found at least 1 microsaccade (a ‘miss’) in the 200 ms time-window surrounding any given time point. Nonetheless, because microsaccades occur only about once or twice every second, both HR and FAR at individual (millisecond) time points will be high (above 0.999). But because HR rose even higher than FAR after stimulus presentation, we found positive values of *d’_o_*.

To extract a single oculomotor sensitivity measure from an entire rate time course for a given condition, we defined a value *o’*, the maximum of the cumulative sum of *d’_o_* values across time (within 200 to 550 ms post-stimulus). *o’* is unaffected by rate rebounds following inhibition, which result in negative *d’_o_*. Pairwise differences in *o’* (across payoff conditions) were tested with bootstrapping, similar to perceptual *d’* as described above.

In addition to the pre-registered analyses reported thus far, we conducted two exploratory analyses. First, to simplify the comparison of microsaccade rates across conditions (without relying on hundreds of noisy tests at many individual time points), we computed the microsaccade rates integrated across two time windows: for the baseline microsaccade rate on target absent trials, we used the time window 0 to 500 ms. For target present trials, we used the time window within which the microsaccade rate on target present trials was significantly lower than the rate on target absent trials, for both bias conditions (bootstrapped FDR-corrected p<0.05). This is the *time window of significant oculomotor freezing* (see results).

Second, compared to our previous studies, we found that baseline microsaccade rates were lower on average and more variable across, which complicates comparing rates by taking simple differences (liberal-conservative) between conditions. We therefore computed *modulation indices* that are more robust to variation across observers in overall microsaccade rates: (A – B) / (A + B), where A and B refer to a measure in specific conditions (e.g., microsaccade rate on conservative vs liberal trials; or report-present vs report-absent trials). This index ranges from −1 to 1, where positive values indicate higher microsaccades rates in A as compared to B, and negative values indicate the opposite.

Finally, we supplement our pairwise tests with Bayes Factors (BFs), which quantify strength of evidence. In this context, a BF is the ratio of the probability of the data under the alternate hypothesis (that two conditions differ), relative to the probability of the data under the null hypothesis (that there is no difference) (Rouder et al. 2009, 2012). As an example, a BF of 10 indicates that the data are ten times more likely under the alternate hypothesis than the null hypothesis. Typically, BFs between 1 and 3 are regarded as weak evidence for the alternate hypothesis, BFs between 3 and 10 as substantial evidence, and BFs between 10 and 100 as strong evidence (Kass and Raftery 1995). Conversely, BFs between than 1/3 and 1/10 are considered substantial evidence for the null hypothesis, etc. We computed BFs for pairwise t-tests and two-way repeated measures ANOVAs using the bayesFactor toolbox by Bart Krekelberg (https://github.com/klabhub/bayesFactor: DOI: 10.5281/zenodo.4394422).

## RESULTS

### Explicit perceptual reports: Bias manipulations affect decision criteria but not sensitivity

On each trial, observers reported the presence or absence of a brief Gabor stimulus with a luminance contrast that had been set to their individual detection threshold. The time of the target’s onset was unpredictable, but the end of each trial was indicated by a beep 500 ms after the time of (potential) target appearance. The observers’ goal was to win “points” that were converted to bonus monetary payments. Correct responses (hits and correct rejections) gained points and incorrect responses (misses and false alarms) lost points.

In Experiment 1, we introduced asymmetric monetary payoffs to manipulate decision bias. In the liberal condition, rewards were three times greater for hits than correct rejections, and penalties were three times greater for misses than false alarms. This reward structure places the optimal criterion at the level of sensory evidence that is three times as likely to be observed when the target is absent than present. Thus, the optimal likelihood ratio *β_opt_* = 1/3). In the conservative condition, rewards were three times greater for correct rejections than hits, and penalties were three times greater for false alarms than misses. That makes *β_opt_* = 3. The reward structure varied across blocks of trials and was known to the participant in advance. Feedback at the end of each trial indicated the reward magnitude.

In Experiment 2, we manipulated probability that a target would appear, and informed observers of that probability on each trial. In the liberal condition, there was a 75% chance that a target would appear (3x likelier to be present than absent), which lowered the optimal criterion such that *β_opt_* = 1/3. In the conservative condition, there was a 25% chance that a target would appear, raising the optimal criterion such that *β_opt_* = 3 (as in Experiment 1). These trial types were randomly intermingled within blocks, but a cue in the form of colored dots presented at the start of each trial informed the participant of the target probability. Payoffs on target-presence and target-absent trials were of equal magnitude.

In both experiments, the bias manipulation strongly affected explicit perceptual reports of target presence. The mean hit and false alarm rates, their mean differences between bias conditions, and the 95% confidence interval (CI) of those differences, are listed in **Table 2**. Hit rates and false alarm rates were much lower in the conservative than liberal condition, indicating that participants were less willing to report seeing a target when the potential payoffs were greater on target absent trials (Experiment 1), and when target presence was unlikely (Experiment 2). Response times are plotted in **Supplementary Figure 1** (https://osf.io/t9by7/).

**Table 2:** Explicit reports in each condition of each experiment. The first two columns list the across-subject mean values, with the standard error in parentheses. The column labeled “Diff” is the average (and SEM) difference: liberal – conservative. The final column is the 95% bootstrapped confidence interval (CI) of the difference. When a CI excludes 0, we conclude there is a significant effect of the bias condition. *d’ and β* are sensitivity and bias measures assuming unequal variance of sensory evidence on target-present and target-absent trials (see text).

|  |  | Conservative | Liberal | Diff | Diff 95% CI |
| --- | --- | --- | --- | --- | --- |
| Hit rate | Expt 1 | 0.57 (0.02) | 0.79 (0.01) | 0.22 (0.03) | [0.18 0.27] |
|  | Expt 2 | 0.41 (0.02) | 0.73 (0.04) | 0.32 (0.05) | [0.22 0.40] |
| False alarm rate | Expt 1 | 0.06 (0.01) | 0.38 (0.04) | 0.31 (0.04) | [0.24 0.40] |
|  | Expt 2 | 0.07 (0.01) | 0.45 (0.07) | 0.38 (0.07) | [0.27 0.52] |
| $d'$ | Expt 1 | 1.95 (0.12) | 1.90 (0.10) | -0.05 (0.08) | [-0.22 0.09] |
|  | Expt 2 | 1.12 (0.09) | 1.27 (0.07) | 0.15 (0.11) | [-0.02 0.40] |
| $\beta$ | Expt 1 | 2.32 (0.38) | 0.42 (0.02) | -1.89 (0.39) | [-2.72 -1.29] |
|  | Expt 2 | 1.80 (0.09) | 0.59 (0.06) | -1.21 (0.11) | [-1.39 -0.99] |

To interpret these psychophysical data, we adopt the classic signal detection model: the participant reports target presence if the magnitude of sensory evidence *E* exceeds a criterion level *c*. The variances of *E* on target-absent and target-present trials are often unequal, and can be estimated with a receiver operating characteristic (ROC) graph (Swets et al. 1961). The ROC in **Figure 1a** plots false alarm rates vs hit rates, each z-transformed through the inverse normal cumulative distribution function. For each participant, one line connects their points for the liberal (blue) and conservative (red) conditions. If the distributions of sensory evidence have equal variance, then these lines should have slopes equal to 1 (illustrated with thick diagonal black lines). The empirical slopes are consistently shallower: in Experiment 1, the mean slope was 0.53 (95% CI = [0.46 0.60]), and in Experiment 2 it was 0.60 (95% CI = [0.50 0.69]). Assuming that the target-absent distributions have standard deviations (SDs) equal to 1, the SD of the target-present distributions are equal to the inverse of the ROC slopes: 1.90 in Experiment 1 and 1.66 in Experiment 2. These best-fitting signal detection models are shown in **Figure 1b**, with the mean criteria (computed directly from false alarm rates) as vertical blue and red lines.

**Figure 1:**
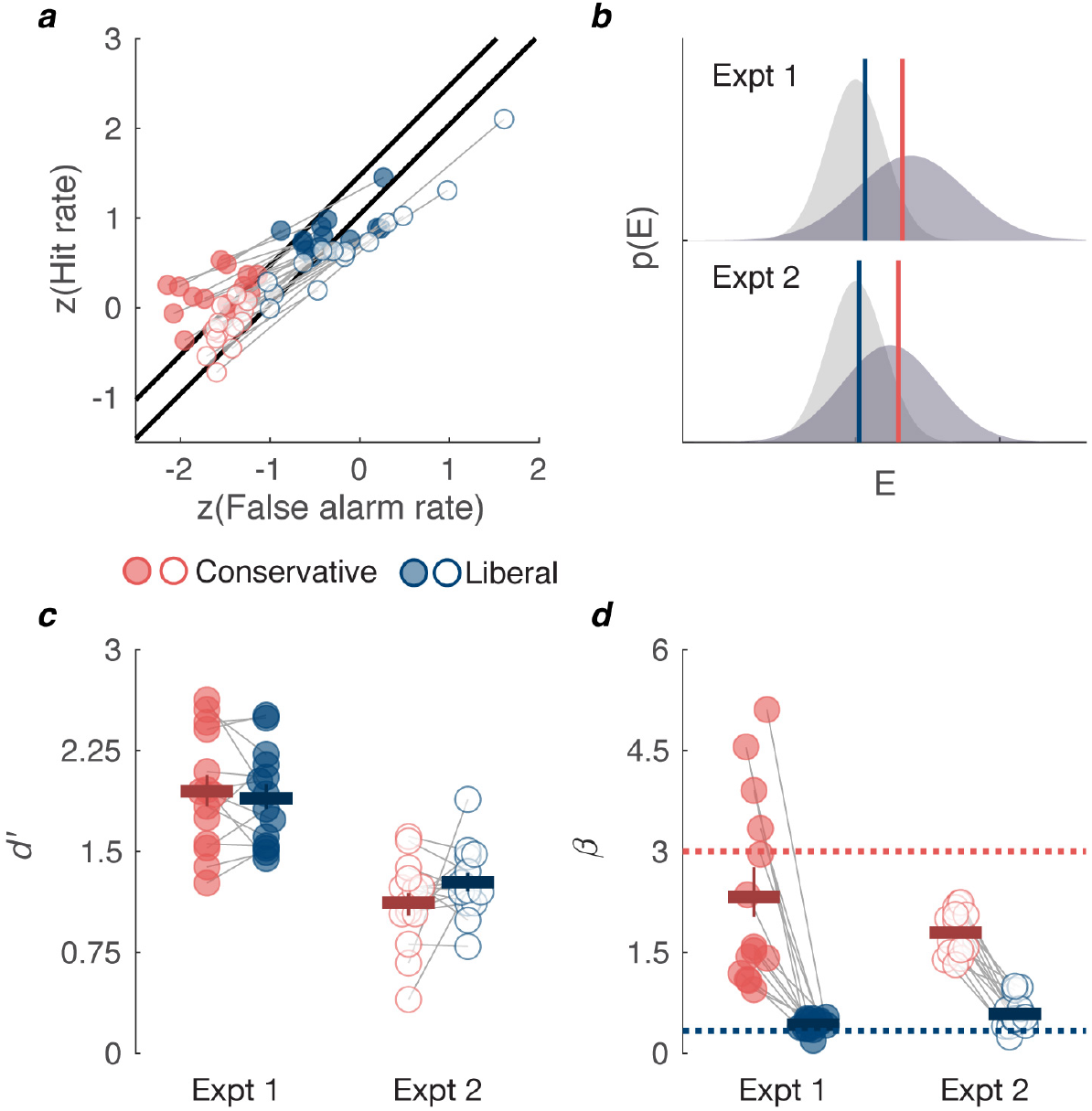
Bias manipulations affect explicit perceptual reports. **(*a*)** The receiver operating characteristic (ROC) showing individual z-transformed hit and false alarm rates. The two black lines with slope 1 are the predictions of an equal-variance model for each experiment (Expt. 1 is the upper black line). The data have slopes consistently less than one, suggesting that the distribution of sensory evidence has higher variance when the target is present rather than absent. **(*b*)** Signal detection models that account for the empirical hit and false alarm rates. These show probability distributions of sensory evidence *E* on target-absent trials (light gray distributions) and target-present trials (darker distributions). The standard deviations of the target-present distributions were derived from the average ROC slopes in panel *a*. The blue and red vertical lines are the mean empirical criteria (computed from false alarm rates) in the liberal and conservative conditions, respectively. **(*c*)** Individual participants’ detection sensitivity *d’*, assuming that the sensory evidence distributions have unequal variance as modeled in panel *b*. Experiment 1 is in filled circles, Experiment 2 in open circles. Thin gray lines connect points from the same participant. The horizontal positions of individual data points are jittered to avoid total overlap, but points from the same participant have the same relative jitter. The horizontal lines represent the means, with error bars spanning the 68% bootstrapped confidence interval (approx. ±1SEM). **(*d*)** Individual participants’ decision bias computed as *β* for each participant, again assuming unequal variance. Format as in panel *c*. Horizontal dotted lines are the optimal *β* for each condition (dark blue = liberal; light red = conservative).

Using these estimated variances, we computed *d’*, a measure of sensitivity (**Figure 1c**), and *β*, a measure of bias (**Figure 1d**). *d’* is the distance between the mean *E* (sensory evidence) on target-present trials and the mean *E* on target-absent trials. *β* is the likelihood ratio of target presence to target absence when *E* = *c*. Using the formulas for *β* and d’ (Equations 2 and 7) that typically assume equal variance, we substituted the best-fitting SDs into the probability and cumulative density functions. Statistics for both measures are reported in **Table 2.** *d’* did not significantly differ between the liberal and conservative conditions (CIs include 0), but *β* was significantly higher in the conservative condition, for all participants. The dashed lines in **Figure 1f** are the optimal *β_opt_* in each condition. Most participants did not shift their criteria quite far enough to reach the optimal levels(Kubovy 1977). For the estimates of *d’* and *β* that (incorrectly) assume equal variance on present and absent trials, see **Supplementary Figure 2** (https://osf.io/t9by7/).

In sum, both bias manipulations had large effects on decision criteria for explicit judgements, while sensitivity remained unaffected.

### Microsaccade rates contingent on physical target presence: Bias manipulations do not affect oculomotor freezing

**Figures 2a** and **2b** show the mean microsaccade rates plotted as a function of time relative to target onset. The target, when present, was flashed at time point 0. In both experiments we observed oculomotor freezing on target-present trials (solid lines): the microsaccade rate begins to drop roughly 130-150 ms after stimulus onset, and then returns to baseline 300-400 ms later. The key question is whether microsaccade rates differ between the liberal and conservative bias conditions. The *distinct-criteria hypothesis* predicts no difference. The *shared-criterion hypothesis*, which posits that oculomotor freezing is linked to explicit report decisions, predicts a greater drop in microsaccade rates on target-present trials of the liberal condition, in which the participant reports “present” more often. The data do not support the shared-criterion hypothesis. Although the mean rate in the liberal condition (blue line) dips slightly lower than in the conservative condition (red line), that effect is small and not consistent across participants.

**Figure 2:**
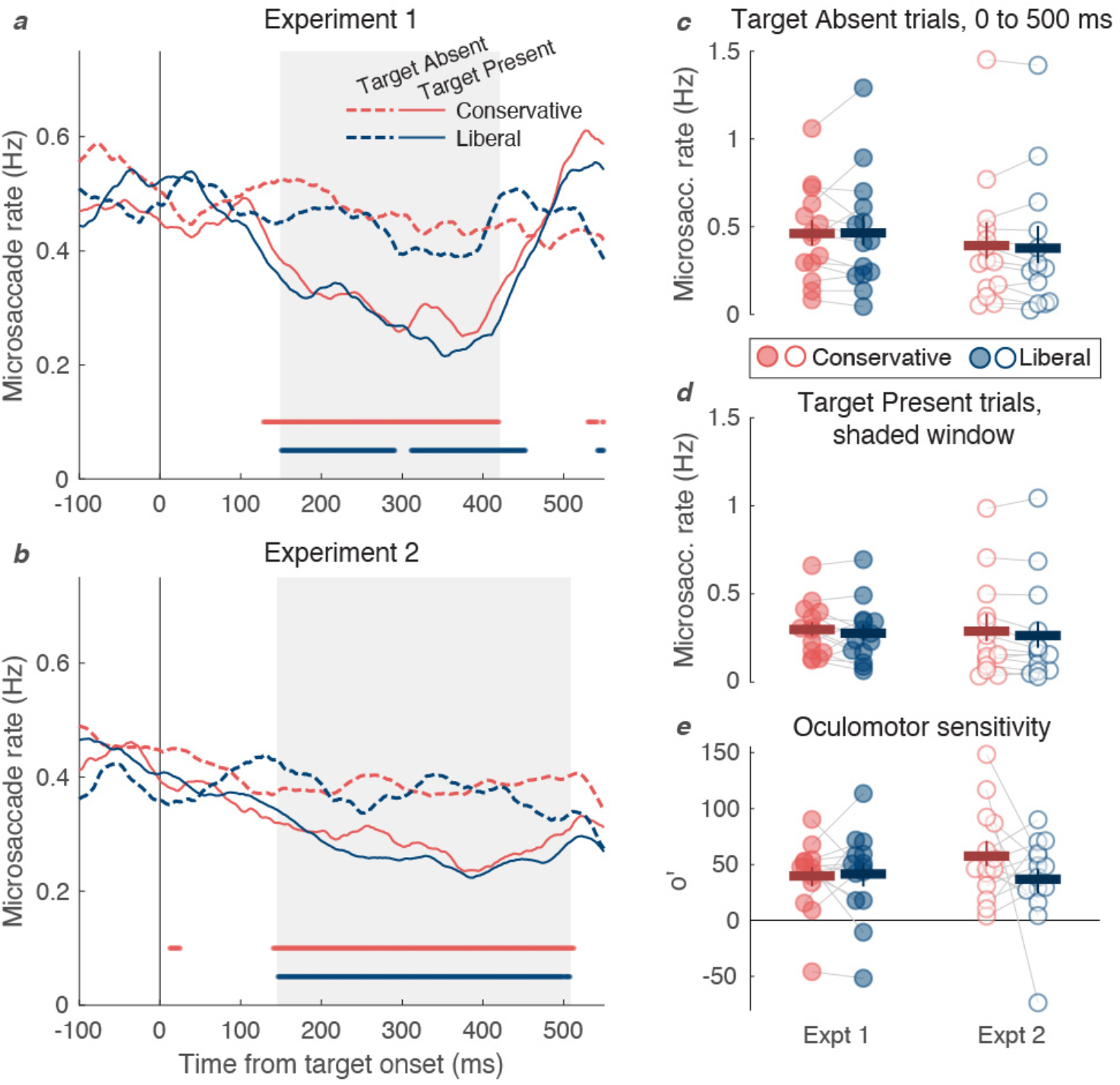
Bias manipulations do not affect overall microsaccade rates on target-present and target-absent trials. **(*a,b*)** Mean microsaccade rates as a function of time relative to target onset in Experiments 1, for target-absent trials (dotted lines) and target-present trials (solid lines). The horizontal lines at the bottom of each plot indicate time points when the rate on target-present trials is significantly different from the rate on target-absent trials (corrected p<0.05). The gray region of the background indicates the time window when the rate was significantly reduced on target-present trials in both conditions. **(*c*)** Mean microsaccade rates on target-absent trials in the time window between 0 and 500 ms. Format as in Figure 1a. **(*d*)** Mean microsaccade rates on target-present trials in the time windows with significant inhibition in both conditions (shaded portions in panels a and b). There are no significant effects of bias condition. **(*e*)** Oculomotor sensitivity (*o’*), a measure of the difference in microsaccade rates between target-present and target-absent trials over the entire interval 0 to 500 ms. There are no significant effects of bias condition.

To simplify this analysis and maximize power, we integrated microsaccades over two key time windows: 0 to 500 ms for target-absent trials and the window of significant oculomotor freezing for target-present trials (shaded windows in **Figures 2a** and **2b**; see **Methods**). In Experiment 1, the window of significant freezing was from 149 ms to 421 ms, and in Experiment 2 it was 145 to 509 ms. As shown in **Figures 2c** and **2d**, there were no reliable effects of bias condition on the mean microsaccade rates in these time windows. We evaluated the effects both as mean differences (L – C, where L is the rate on liberal trials and C on conservative trials) and as modulation indices [(L – C)/(L + C)] that adjust for individual differences in overall microsaccade rate. With one exception, none of those effects were significant: 95% CIs include 0, and Bayes Factors (BFs) support the null hypothesis at least 2:1 (BFs<0.5). The one exception is for target-absent trials in Experiment 2: when the effect is expressed as a modulation index, the baseline microsaccade rate was slightly but significantly lower on liberal than conservative trials (mean index = −0.09, 95% CI = [−0.17 - 0.02]; BF=1.35. The mean difference was only −0.02 Hz (95% CI = [−0.04 0.03]; BF=0.37).

To combine across experiments, we entered these data into linear mixed effects models (LMEs), with fixed effects for condition, experiment, and their interaction, as well as random effects for participant. We fit one such model for the target-absent trials and another for the target-present trials. The effect of condition was negligible in both analyses (0.006 and 0.02 Hz, respectively), and not significant (both p>0.10; both BF<0.5). There were no effects of experiment nor interaction between experiment and condition (all p>0.5).

We also computed oculomotor sensitivity (*o’*) as a measure of the strength of oculomotor freezing (White and Rolfs 2016) (**Figure 2e**), comparable to *d’*. In both experiments, *o’* did not differ significantly between bias conditions: 95% CIs were far from excluding 0 and Bayes Factors supported the null hypothesis (all BFs<0.5). If anything, the effect in Experiment 2 (conservative > liberal) went in the direction opposite predicted by the shared criterion hypothesis, but was not significant (mean modulation index = −0.32, 95% CI = [−1.55 0.11]). An LME combining across experiments found no effect of condition (*p*=0.32, 95% CI = [−28.4 9.5]; BF=0.30) and no main effect of experiment nor interaction (both *p*>0.2).

Altogether, the microsaccade rates in this first analysis are consistent with the distinct-criteria hypothesis: oculomotor freezing is independent of bias manipulations that affect explicit perceptual reports. Next, we sorted the data further by the participant’s report on each trial. Based on our prior study (White and Rolfs 2016), we predicted more oculomotor freezing on trials when the participant reports seeing a stimulus than when they don’t, but the magnitude of that effect may depend on the bias condition.

### Microsaccade rates contingent on explicit perceptual reports: Oculomotor freezing is stronger in conservative than liberal bias conditions

When we analyze trials separately according to whether the participant reported target presence or absence, the shared-criterion hypothesis predicts no effect of bias condition. The observer’s ultimate decision is the same on liberal hit trials as on conservative hit trials, so the prevalence of oculomotor freezing should be the same. In contrast, the distinct-criteria hypothesis predicts an effect of bias condition: when considering only trials in which the observer reports target presence (hits and false alarms), microsaccade rates should be lower in the conservative condition than liberal condition. This is because in the conservative condition, the sensory evidence must be stronger for the participant to report presence, and therefore it is also likely to trigger oculomotor freezing. In the liberal condition, some explicit reports of target presence are guesses with low sensory evidence, which will not exceed the criterion for oculomotor freezing, so microsaccade rates should be higher.

**Figure 3a** plots the mean microsaccade rates on target absent trials, separated by bias condition and the participant’s explicit report of whether a target was present or not (correct reject trials in dark lines, false alarm trials in bright lines). In a prior study (White and Rolfs 2016), we found that microsaccade rates were lower on false alarm than correct reject trials, consistent with the notion that a spurious sensory signal triggered both an explicit false alarm and oculomotor freezing. The distinct-criteria hypothesis predicts that effect (the relative inhibition of microsaccades on false alarm trials) should be weakened in the liberal condition, when many false alarms are guesses without a sensory signal strong enough to inhibit microsaccades.

**Figure 3:**
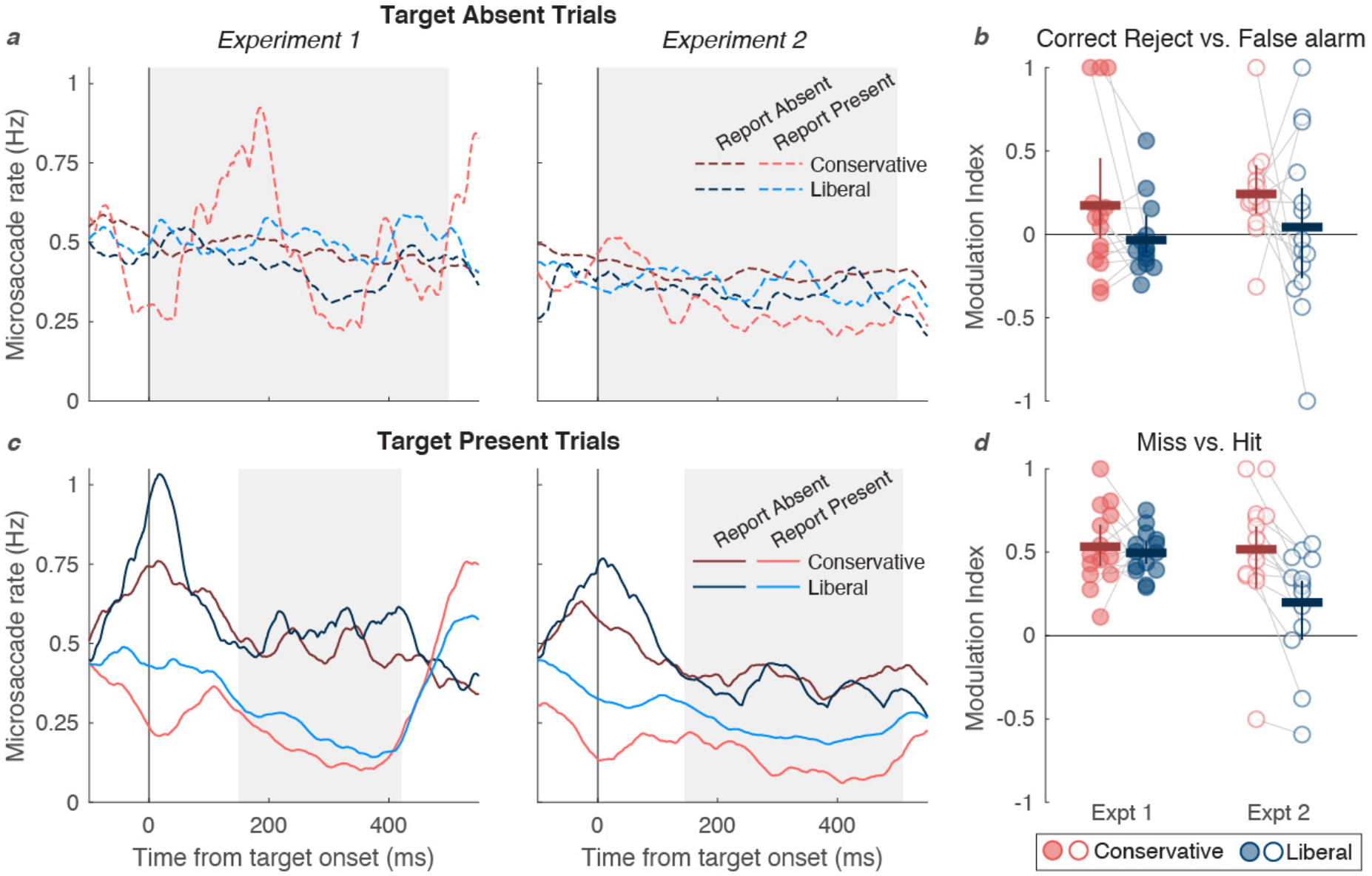
Microsaccade rate signatures as a function of bias condition and explicit report outcome. **(*a*)** Mean rates as a function of time on target absent trials, separated by bias condition and by whether the participant reported target absent (correct reject trials, dark lines) or target present (false alarm trials, bright lines). Note there are very few false alarm trials in the conservative condition (bright red lines) **(*b*)** The mean modulation indices comparing microsaccade rates on correct reject trials and false alarm trials, integrated over 0 to 500 ms (shaded interval in panel *a*). Format as in Figure 1a, except the error bars are bootstrapped 95% confidence intervals to highlight significant deviations from zero. The overall effect of perceptual report is significant, and marginally higher on conservative than liberal trials. **(*c*)** Mean microsaccade rates on target present trials, separated by bias condition and by whether the participant reported target absent (miss trials, dark lines) or reported target present (hit trials, bright lines). **(*d*)** The mean indices comparing microsaccade rates on miss and hit trials, integrated over the intervals with significant stimulus-induced inhibition (shaded in panel c). Microsaccade rates are significantly lower on hit than miss trials, and that effect is significantly larger on conservative than liberal trials.

To test these predictions, we integrated microsaccade rates over the 0 to 500 ms time window (shaded region) and then computed the effect of explicit report as a modulation index: (CR – FR) / (CR + FR), where CR is the microsaccade rate on correct reject trials and FR is the microsaccade rate on false alarm trials. The occurrence of oculomotor freezing on false alarm trials predicts a positive index. In addition, the distinct-criteria hypothesis predicts a larger index in the conservative compared to the liberal condition. The mean indices are plotted in **Figure 3b** and listed in Table **2** with 95% CIs and BFs. Only in the conservative condition of Experiment 2 was the effect of report significant. According to a linear mixed effects model that combined experiments, there was a small but significant difference between microsaccade rates on correct reject vs. false alarm trials (mean index = 0.11, CI = [0.004 0.209], p=0.04, BF=1.4), a marginal effect of bias condition (index 0.2 larger in the conservative condition, CI = [0.002 0.41], p=0.053; BF=1.28), and no effect of experiment nor interaction (both p>0.4, BF<0.25). All told, the data in **Figure 3b** support our previous finding that false alarms are associated with inhibition of microsaccades, and are consistent with the distinct-criteria hypothesis. However, this analysis is limited due to the small number of false alarm trials in the conservative condition (on average across participants, only 20 trials in Experiment 1 and 70 in Experiment 2). The target-present trials provide supporting evidence.

**Figure 3c** plots microsaccade rates on target-present trials. These traces diverge around stimulus onset (i.e., 0 ms) due to the reductive effect of microsaccades on perceptual sensitivity (Rolfs 2009; Scholes et al. 2018; White and Rolfs 2016; Zuber et al. 1964): a microsaccade that occurs close in time to the stimulus onset can make the participant miss the stimulus, thus miss trials are associated with a peak in the microsaccade rate near time 0. That peak is especially large in the liberal condition, when misses are less frequent and require a definite lack of target evidence. Conversely, hits are associated with fewer microsaccades near the time of stimulus onset, and thus there is a dip in microsaccade rate on hit trials. That dip is larger in the conservative condition, when hits require high certainty and would otherwise be turned to misses by microsaccades. We previously confirmed that the drop in microsaccade rates on hit trials ~150-400 ms post-stimulus is not an artifact of the divergent dips and peaks observed around 0 ms due to saccadic suppression of perception (White and Rolfs 2016).

Our current research question focuses on the later time period, starting roughly 150 ms post-stimulus, when stimulus detection is associated with inhibition of microsaccades. We tested whether that effect of perceptual report (misses vs hits) is equal in the two bias conditions.

The distinct-criteria hypothesis predicts greater inhibition on hit trials of the conservative condition, because conservative hits are “purer” (i.e., they contain fewer lucky guesses) and require a strong sensory signal that is also likely to trigger oculomotor freezing.

Indeed, the microsaccade rate dips lower on hit trials of the conservative condition (**Figure 3c**, light red lines) than of the liberal condition (light blue lines). To summarize these effects, we integrated microsaccade rates over the time window with significant inhibition (shaded windows in **Figures 2a**, **2b**, and **3b**). For each bias condition we then computed the effect of explicit detection as a modulation index: (M-H)/(M+H), where M is the microsaccade rate on miss trials and H is the rate on hit trials. The effect of explicit detection was significant (95% CI of the index excludes 0) in all conditions except the liberal condition of Experiment 2 (see **Table 3**). According to a linear mixed effects model, that modulation index was significantly larger in the conservative than liberal condition (by 0.18 on average, CI = [0.08 0.14], p=0.0004; BF=34.0). This is strong evidence that correct reports of target presence in the conservative condition are associated with stronger inhibition of microsaccades than in the liberal condition. The effect of bias condition was also larger in Experiment 2 than Experiment 1 (interaction between condition and experiment, p=0.004; BF=6.32).

**Table 3:** Effects of perceptual report on microsaccade rates, expressed as modulation indices. The columns labeled “Modulation index” contain the mean, with standard error across participants in parentheses. The 95% CIs are derived from bootstrapping.

|  |  | Correct reject - False alarm |  |  | Miss - Hit |  |  |
| --- | --- | --- | --- | --- | --- | --- | --- |
|  | Condition | Modulation index | Index 95% CI | BF | Modulation index | Index 95% CI | BF |
| **Expt. 1** | Conservative | 0.17 (0.13) | [-0.039 0.437] | 0.58 | 0.53 (0.06) | [0.429 0.657] | $1.4 \times 10^4$ |
| | Liberal | -0.03 (0.06) | [-0.121 0.116] | 0.41 | 0.50 (0.04) | [0.433 0.566] | $2.6 \times 10^6$ |
| **Expt. 2** | Conservative | 0.24 (0.08) | [0.128 0.403] | 7.47 | 0.52 (0.10) | [0.248 0.671] | 197.6 |
|  | Liberal | 0.04 (0.14) | [-0.226 0.319] | 0.27 | 0.20 (0.09) | [-0.008 0.333] | 1.68 |

These data consistently support the distinct-criteria hypothesis: oculomotor freezing is triggered when a sensory signal crosses a threshold that is independent of the participant’s decision bias. The sensory signal is more likely to have crossed that oculomotor threshold on hit trials of the conservative condition, when the criterion for explicit report is higher, than on hit trials of the liberal condition. Thus, when the participant was induced to adopt a more conservative decision bias, explicit detection of the stimulus was associated with more pronounced oculomotor freezing.

## DISCUSSION

Detecting potentially relevant stimuli in the environment is a fundamental task of perceptual systems. Our data suggest that although sensory input is continuous and noisy, the brain switches into a qualitatively different state when there is sufficient evidence that a target is present. Passing this threshold gives rise to a conscious percept *and* an involuntary pause of saccadic eye movements (that is, oculomotor freezing). A pause in microsaccades can be considered the oculomotor system’s “report” that it detected a stimulus. The participant’s decision to respond voluntarily to the stimulus — for instance, by pressing a button — depends on the conscious percept as well as potential rewards and expectations.

Visual stimulus detection therefore has three consequences that are of interest to the present investigation: a conscious percept, a decision to report stimulus presence, and oculomotor freezing. It is crucial that we understand how those three consequences relate in terms of neural and cognitive mechanisms. Provided oculomotor freezing is indeed a proxy for conscious perception (as we argue below), researchers would be equipped with a “no-report” paradigm to investigate the neural correlates of consciousness (Tsuchiya et al. 2015) without interference from explicit cognitive tasks.

In five independent experiments across this study and a previous one (White and Rolfs 2016), we consistently found that explicit reports and oculomotor freezing covary: the eyes only freeze in response to stimuli that the person reports seeing. To explain that covariation, here we manipulated the likelihood of explicit reports of stimulus presence. When rewards and penalties were greater on target-present than target-absent trials (Experiment 1), or when the target probability was known to be high (Experiment 2), participants adopted a liberal decision criterion, reporting target presence much more often than in the opposite (conservative) conditions (**Figure 1**).

In contrast, the magnitude of the drop in microsaccade rates just after stimulus onset showed little to no effect of our bias manipulations (**Figure 2**). We need not rely only on that null result, however, because we also found effects of the bias condition when splitting the trials according to the explicit report (**Figure 3**). The difference in microsaccade rates between hit and miss trials, which indexes the link between explicit reports and oculomotor freezing, was larger in the conservative than the liberal condition. Our interpretation is that when participants make conservative decisions, they only report sensations that are strong enough to also trigger oculomotor freezing. In contrast, when participants make liberal decisions, they often make strategic guesses that a stimulus was present, even when the sensory signal was weak and oculomotor freezing was not triggered.

We therefore reject the shared-criterion hypothesis and support the distinct-criteria hypothesis (described in the Introduction). The criteria in question specify the magnitude of sensory evidence required to trigger a response. There is one criterion for explicitly reporting stimulus presence, and it can be shifted to maximize rewards. There is also a distinct criterion for inhibiting eye movements, which is relatively stable and not affected by shifts of decision criterion.

This conclusion is consistent with studies that predicted individual perceptual contrast thresholds based on microsaccade patterns that were measured while the participant did not explicitly respond to the stimuli (Bonneh et al. 2015; Denniss et al. 2018; Scholes et al. 2015). These studies show that oculomotor freezing is not related to response preparation. However, participants in those studies either had to silently count the stimuli (Bonneh et al. 2015), or prepare to respond on a random subset of trials (Denniss et al. 2018; Scholes et al. 2015), so they were likely making covert decisions about each stimulus. Therefore, decision-making processes could have contributed to oculomotor freezing in those data. Our data help isolate the link between perception and oculomotor freezing.

A key feature of our theory is that oculomotor freezing is all-or-none, not graded. In a prior study (White and Rolfs 2016), we varied the visibility of a target grating by varying its luminance contrast, or by adapting the observer to the same or different orientation. Considering all target-present trials, the degree of oculomotor freezing scaled with explicit *d’*. However, when considering only hit trials, oculomotor freezing was equivalently across all contrast levels and adaptation states. Intense stimuli had no effect on eye movements if the observer missed them, and faint stimuli were accompanied by full-fledged inhibition provided they were detected. We found similar patterns in the new data reported above, providing consistent support that oculomotor freezing is a discrete all-or-none reflex that occurs if and only if a stimulus is consciously detected. Such a model is reminiscent of “high threshold theory,” which has been largely discredited (Swets 1961). Standard signal detection theory, which has been more successful, assumes no threshold for detection other than the observer’s flexible decision criterion. In that regard, the data presented here are not fully consistent with standard signal detection theory.

Our data are, at least in part, consistent with the “global neuronal workspace theory” of consciousness. It proposes that a stimulus becomes reportable when “ignites” sustained neural communication across the brain (Mashour et al. 2020). Recent electrophysiological data suggest that ignition occurs when activity in frontal cortex, not sensory cortex, reaches a threshold, roughly 200 ms after stimulus onset (Van Vugt et al. 2018). We might speculate that such an ‘ignition’ is related to oculomotor freezing, but it occurs later than the initial drop in microsaccade rates.

For now, we consider two possibilities for how oculomotor freezing relates to the conscious experience of the stimulus that triggers it. Both are compatible with the distinct-criteria hypothesis described in the Introduction. (1) Oculomotor freezing and conscious perception are tightly coupled, because they are both triggered when the same sensory signal crosses the same threshold. That threshold is not affected by bias manipulations, unlike the criterion for explicit reports. (2) Oculomotor freezing and conscious perception can be dissociated, because the threshold for oculomotor freezing is stable but the threshold for conscious perception is affected by bias manipulations, along with the decision criterion. While our data reveal that explicit reports and oculomotor freezing have distinct criteria, they are consistent with both possibilities regarding conscious perception.

We nonetheless favor the first possibility: oculomotor freezing and conscious perception are coupled. This hypothesis must also assume that the bias manipulations (payoffs and probability cues) affect decisions at a post-perceptual stage. Specifically, in the liberal conditions, observers reported “yes” more often because doing so maximized rewards, not because they actually *saw* the target more often. That is why the difference in microsaccade rates between hit and correct reject trials is weaker in the liberal than conservative condition. The implication is that oculomotor freezing provides an implicit index of conscious perception that is free of bias.

However, this conclusion fails if the bias manipulations do affect conscious perception (i.e., the second possibility). There is some neurophysiological evidence that expectations, as manipulated by the probability cues in Experiment 2, can affect sensory processing (De Lange et al. 2018; Pajani et al. 2015). One theory is that expecting a stimulus evokes a “template” in neural populations that prefer the expected features (Kok et al. 2014, 2017). In contrast, one fMRI study concluded that payoff and probability manipulations recruit frontal and parietal brain regions involved in decision-making to shift the starting point of evidence accumulation, similar to a criterion shift (Mulder et al. 2012). The existing behavioral evidence is also ambiguous. One study argued that expectation improves detection by elevating the baseline of “signal-selective units” (Wyart et al. 2012). Another found that probability cues presented *after* the stimulus had similar effects as cues presented *before* the stimulus, in favor of a post-perceptual criterion shift (Bang and Rahnev 2017). It remains a matter of discussion, therefore, whether expectations affect conscious perception or decision processes (Press et al. 2020; Rungratsameetaweemana et al. 2018; Rungratsameetaweemana and Serences 2019; Summerfield and Egner 2016). The simplest model that explains our data assume that they affect decision processes only.

We must also note that the results favoring the distinct-criteria hypothesis are clearer in Experiment 2 (which manipulated expectations) than Experiment 1 (which manipulated rewards). Indeed, other researchers have found that probability manipulations have stronger effects on perceptual decisions than reward manipulations do (Leite and Ratcliff 2011; Mulder et al. 2012; Simen et al. 2009). In our case there are several possible explanations: first, there were greater individual differences in explicit report criteria in Experiment 1 (**Figure 1d**), perhaps due to variable interpretations of, or value placed in, the rewards. Such individual differences may have added noise to the microsaccade data as well. Second, overall *d’* levels were higher in Experiment 1 than 2 (**Figure 1c**). The bias manipulations are likely to have greater effects when the target is difficult to detect. Third, it may be that expected rewards affect decisions at a post-perceptual stage, whereas expectations affect perception, as discussed above. In that case, the explicit reports in Experiment 1 were less driven by the sensory signal, thus showing a less clear relationship with oculomotor freezing. In contrast, Experiment 2 could be explained by a model in which expecting a stimulus lowers the sensory threshold for conscious perception, but does not affect the threshold for oculomotor freezing. While this model would explain the smaller difference in microsaccade rates between hit and miss trials in the liberal condition (**Figure 3d**), it is comparably complicated.

We may also consider the probability cues of Experiment 2 in light of the “predictive coding” framework (De Lange et al. 2018). A target in the “conservative” condition is unexpected, and thus should produce a larger prediction error. If oculomotor freezing is a “surprise” response, then we would have predicted a larger drop in microsaccades in target-present trials of the conservative condition than the liberal condition. We did observe that, but only on hit trials (**Figure 3c**). The predictive coding framework may therefore help explain oculomotor freezing.

Altogether, the most parsimonious explanation for our results is that oculomotor freezing and conscious detection share a common sensory threshold. This threshold is distinct from the decision criterion to report a stimulus, which is shifted by weighted payoffs and expectations. The alternate explanation, that the threshold for conscious detection can diverge from the threshold for oculomotor freezing, is more complicated. It must either postulate an additional free parameter, for a total of three sensory thresholds/criteria, or it must assume that the decision criterion is also the threshold for perception and thus bias manipulations truly affect perception. To the extent that the more parsimonious explanation stands, oculomotor freezing provides a valuable tool to measure conscious perception free of the influence of decision bias, and without requiring explicit reports.

## Supplementary Data

Two supplementary data figures can be viewed here: https://osf.io/t9by7/.

## Acknowledgments

We are grateful to Michael Grubb, Jan-Nikolas Klanke and Dobromir Rahnev for comments on this manuscript, to Jan-Nikolas Klanke for data collection, and to Richard Schweitzer for technical assistance.

## Author contributions

Conceptualization: AW, JM, MR

Methodology: AW, JM, MR

Investigation: AW, MR

Software: AW, MR

Resources: MR

Formal analysis: AW

Visualization: AW

Supervision: MR

Writing—original draft: AW

Writing—review & editing: AW, MR

