## Supplemental Figures for "Oculomotor freezing indicates conscious detection free of decision bias"

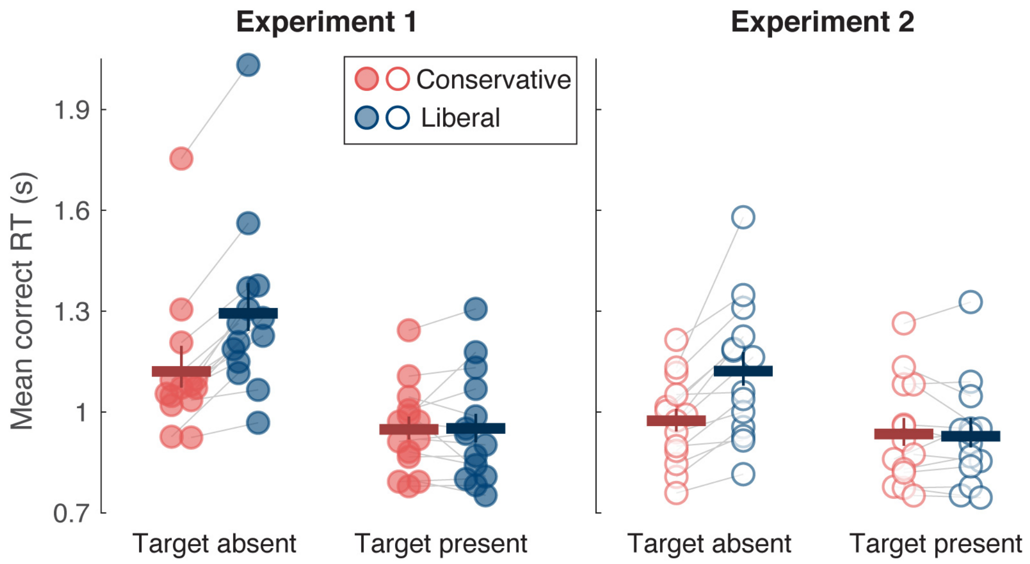


**Figure S1:** Geometric mean of correct response times (RTs) on target-absent and target present trials, separately in each experiment. RTs were generally slower on target-absent than target-present trials. Within target-absent trials (correct rejections), responses were slower in the liberal than conservative conditions (Experiment 1: mean difference = 0.173 s, CI = [0.130 0.212], BF = 8551; Experiment 2: mean difference = 0.148 s, CI = [0.105 0.229], BF = 125). There was no effect of bias condition on correct responses times on target-present trials (hits) in either experiment (mean differences = 0.003 and -0.007 s, BFs<0.3). These experiments were not optimized for drawing inferences based on RTs, because the participants were not allowed to respond until a beep that occurred 500 ms after the time of (potential) target onset. Nonetheless, one interpretation of the pattern on target-absent trials is that in the liberal condition, observers only report "target absent" when they took some time to think about what they saw and are confident. Therefore, they are slower than in the conservative condition when reporting "target absent" is the default response.

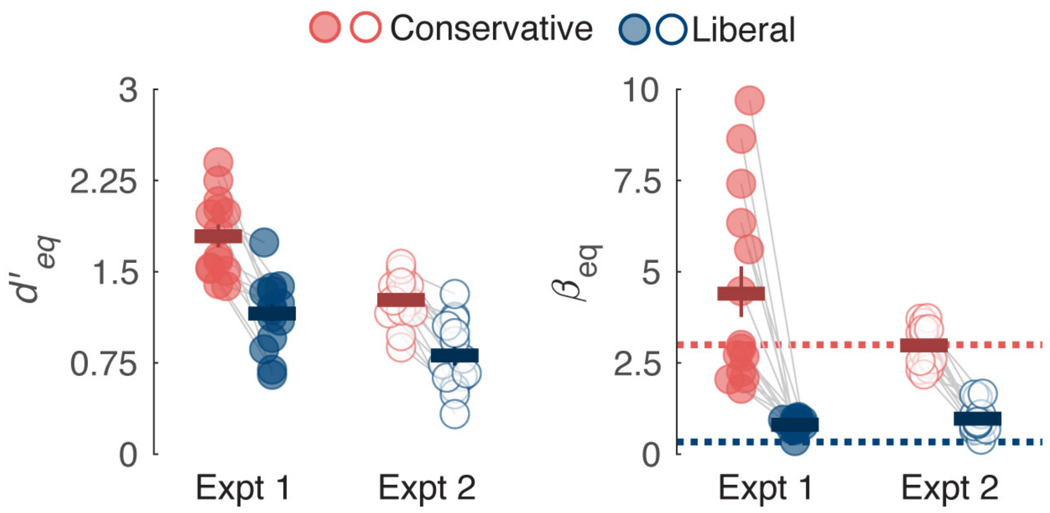


**Figure S2:** Estimates of sensitivity and bias computed assuming equal variance of sensory evidence *E* on target-present and target-absent trials, to yield $d'_{eq}$ and $\beta_{eq}$. Format as in Figure 1c. $d'_{eq}$ was significantly lower in the liberal than conservative condition (mean difference = 0.63 and 0.46 in Experiments 1 and 2 respectively), which seems to violate the tenet of signal detection theory that sensitivity is independent of decision bias. However, as illustrated in the main text and Figure 1a, we can reject that assumption of equal variance. The corrected estimates of sensitivity show no difference between conservative and liberal conditions.
